# Sensory processing sensitivity and axonal microarchitecture: Identifying brain structural characteristics for behavior

**DOI:** 10.1101/2021.11.13.468491

**Authors:** Szabolcs David, Lucy L. Brown, Anneriet M. Heemskerk, Elaine Aron, Alexander Leemans, Arthur Aron

## Abstract

Previously, researchers used functional MRI to identify regional brain activations associated with sensory processing sensitivity (SPS), a proposed normal phenotype trait. To further validate SPS as a behavioral entity, to characterize it anatomically, and to test the usefulness in psychology of methodologies that assess axonal properties, the present study correlated SPS proxy questionnaire scores (adjusted for neuroticism) with diffusion tensor imaging measures. Participants (n=408) from the Young Adult Human Connectome Project that are free of neurologic and psychiatric disorders were investigated. We computed mean diffusivity (MD), radial diffusivity (RD), axial diffusivity (AD) and fractional anisotropy (FA). A voxelwise, exploratory analysis showed that MD and RD correlated positively with SPS proxy scores in the right and left subcallosal and anterior ventral cingulum bundle, and the right forceps minor of the corpus callosum (peak Cohen’s D effect size = 0.269). Further analyses showed correlations throughout the entire right and left ventromedial prefrontal cortex, including the superior longitudinal fasciculus, inferior fronto-occipital fasciculus, uncinate and arcuate fasciculus. These prefrontal regions are generally involved in emotion, reward and social processing. FA was negatively correlated with SPS proxy scores in white matter of the right premotor/motor/somatosensory/supramarginal gyrus regions, which are associated with empathy, theory of mind, primary and secondary somatosensory processing. Region of interest (ROI) analysis, based-on previous fMRI results and Freesurfer atlas-defined areas, showed small effect sizes, (+0.151 to -0.165) in white matter of the precuneus and inferior frontal gyrus. Other ROI effects were found in regions of the dorsal and ventral visual pathways and primary auditory cortex. The results reveal that in a large, diverse group of participants axonal microarchitectural differences can be identified with SPS traits that are subtle and in the range of typical behavior. The results suggest that the heightened sensory processing in people who show SPS may be influenced by the microstructure of white matter in specific neocortical regions. Although previous fMRI studies had identified most of these general neocortical regions, the DTI results put a new focus on brain areas related to attention and cognitive flexibility, empathy, emotion and low-level sensory processing, as in the primary sensory cortex. Psychological trait characterization may benefit from diffusion tensor imaging methodology by identifying influential brain systems for traits.

## 1. Introduction

### 1.1 Sensory processing sensitivity (SPS)

Sensory processing sensitivity (SPS) (Aron and Aron, 1997) is proposed to be a normal phenotype trait observable as a high degree of environmental sensitivity (Pluess, 2015). It is unrelated to sensory processing disorder. Heritability explains about 42% of the variance in twin studies (Assary et al., 2020). The trait is present in about 20% of the population (Lionetti et al., 2018); the estimated number varies from 15% (Kagan, 1997) to 30% (Pluess et al., 2018). A similar proportion is found in many other species, suggesting the presence of two major survival strategies (Wolf et al., 2008): (1) SPS: A high level of attention to environmental stimuli and observing carefully before acting and (2) a normal level of attention to the environment and thus being the first to act. High attention to the environment appears only in a minority of a population because this high sensitivity benefits an individual only if the majority of members of the species lack it (negative frequency dependence).

In humans this increased sensitivity compared to the general population is hypothesized to result from a greater depth of processing of sensory input, e.g. using many cognitive tags (Aron et al., 2012; Aron and Aron, 1997). Lockart et. al. (Lockhart et al., 1976) suggested that memory is enhanced by processing information to deeper cognitive levels by using a letter, word, word in context, or word abstracted from many contexts, a concept important to educational methodology (Leow, 2018). This type of processing, using semantics or many contexts to remember a visual stimulus, is thought to enhance awareness of subtle stimuli and increase emotional responsiveness to stimuli in people with SPS. The emotional responsiveness may in turn act as a motivator for the depth of processing (Baumeister et al., 2007).

Research on SPS is growing, for a review see (Greven et al., 2019). For example, research using fMRI found SPS was associated with significantly greater activation in brain areas involved in higher-order visual processing, which is more evidence for greater depth of processing in SPS compared to the general population. However, there are many open questions, especially about neuroanatomical correlates. There has been no previous research that investigates microstructural changes using diffusion tensor imaging or similar techniques.

SPS does not appear to be a disorder, given the percentages in the population, its presence in many species, and its functionality as a successful survival strategy. In a review comparing fMRI studies of SPS (Acevedo et al., 2018), autism spectrum disorder, schizophrenia, and post-traumatic stress disorder, the authors conclude that SPS engages brain regions differently from these disorders, namely those that are involved in reward processing, memory, physiological homoeostasis, self-other processing, empathy, and awareness. However, SPS can be related to disorders in that it leads to greater susceptibility to environmental influences (Belsky and Pluess, 2009). For example, adults high on SPS scores who report difficult childhoods are more prone to depression, anxiety, and shyness (Aron et al., 2005); however, young children who exhibit SPS and had especially good childhood environments performed well on measures of social and academic competence (Pluess and Belsky, 2010), while those with worse environments fared poorly.

The trait also has been shown to lead to greater positive outcomes following interventions (Nocentini et al., 2018; Pluess and Boniwell, 2015), again suggesting that scores on the measure are correlated with greater susceptibility to environmental influences. For example, a study (Karam et al., 2019) of Syrian refugee children suggests that high-SPS scorers’ previous experience with family trauma (abuse, neglect, etc.) appeared to prepare them to be less affected by war trauma, while those with positive family histories reported more trauma from similar war-related events, again suggesting a survival strategy of noticing more rather than simply needing a positive childhood to adjust well to any circumstance. Differential susceptibility makes the sample in the present study particularly valuable given that participants were screened to avoid those with psychological disorders, that axonal microstructural differences possibly due to a seriously problematic past would not confound our results.

### 1.2 The Young Adult-Human Connectome Project

The dataset of the Young Adult Human Connectome Project (YA-HCP) offered a large group to analyze for a normal psychological variable and any relationship it might have to axonal microarchitectural measures. This open data cohort includes high quality imaging data and an extensive range of data analysis options. There is also extensive psychological testing for each participant.

### 1.3 Proxy scale used to measure SPS

The questionnaire measure for SPS used in previous studies is called the HSP or Highly Sensitive Person Scale (Aron and Aron, 1997). The YA-HCP testing does not include the HSP Scale and it was not feasible to re-contact participants to administer it. However, the YA-HCP dataset does include a substantial number of multi-item self-report personality measures. Thus, it was possible to identify a subset of items in the YA-HCP dataset that could serve as a proxy measure. As described in the Methods, we developed and validated in other groups a 17-item proxy scale. We call this scale that measures SPS in the present report the neuroticism-adjusted residual proxy HSP scale, or Proxy HSP Scale.

### 1.4 Microstructural characteristics: Diffusion Tensor Imaging

Microstructural characteristics associated with SPS would be another strong piece of evidence that it is a significant, reliable psychological trait. Also, identification of brain regional effects contributes to better understanding of SPS functional systems. Thus, we used diffusion tensor imaging (DTI) to identify any possible axonal microstructural characteristics for SPS. We captured measures from mean diffusivity (MD), radial diffusivity (RD), axial diffusivity (AD) and fractional anisotropy (FA). First, we used an exploratory voxelwise analysis for the whole brain. In addition, we used fMRI data from previous studies to carry out a region of interest (ROI) analysis.

### 1.5 Previous fMRI studies

Several fMRI studies helped to validate SPS as a psychological trait affecting sensory processing by finding correlations between the standard self-report measure of SPS and neurophysiological events during a variety of perceptual tasks. These studies provided the locations for the ROI analysis. The first study used a task of perceiving subtle differences in neutral landscapes (Jagiellowicz et al., 2011). When detecting minor (vs major) changes in the landscape, high scores on the standard HSP Scale were associated with greater activation in brain areas involved in higher-order visual processing: left occipitotemporal, bilateral temporal, and medial and posterior parietal regions.

Another fMRI study looked at culturally influenced visual perception (Hedden et al., 2008). The researchers gave 10 European-Americans and 10 East-Asians a visuospatial task that was either context independent (judging the length of a line independent of a surrounding box, the absolute condition, typically harder for Asians) or context dependent (judging the length of a line while paying attention to the box, the relative condition, typically harder for Americans). Each group exhibited greater activation for the culturally non-preferred task in frontal and parietal regions associated with greater effort in attention and working memory. In the two cultural groups, the HSP Scale scores moderated the brain activations such that neither cultural group with high HSP scores showed greater activation on their culturally more difficult task (Aron et al., 2010). The data suggest that the high-SPS participants were processing both the relative and absolute conditions, unaffected by their culture, by paying close attention to details of the stimulus.

Another fMRI study used visual stimuli that were photos of familiar or unfamiliar faces with happy, neutral, or sad expressions (Acevedo et al., 2014). Across all conditions standard HSP scores were associated with increased brain activation of regions involved in attention and action planning (in the cingulate and premotor area (PMA). For happy- and sad-face conditions, SPS was associated with activation of brain regions involved in self-awareness, integration of sensory information, empathy, and action planning (e.g., cingulate, insula, inferior frontal gyrus [IFG], middle temporal gyrus [MTG], and premotor area [PMA]).

Finally, SPS individuals showed substantial differences compared to others in brain activation in response to emotional (versus neutral) images (nonsocial visual International Affective Picture System images; (Acevedo et al., 2017)). Standard HSP scores were associated with neural activations in the temporal/parietal area and areas that process emotional memory, learning, awareness, reflective thinking, and integration of information. There were similar results in the same study for an SPS x Quality of Childhood Parenting (QCP) interaction. For positive stimuli, SPS showed significant correlations with activation in subcortical areas involved in reward processing, self-other integration (insula and IFG), calm (PAG), and satiation (subcallosal AC). These were stronger with increasing QCP. For negative stimuli, the SPS x QCP interaction showed significant activation in the amygdala and prefrontal cortex (PFC) involved in emotion and self-control.

Overall, these fMRI studies show that SPS is associated with greater activation in multiple brain areas when processing subtle visual differences in neutral stimuli as well as stimuli evoking emotion or empathy and personally relevant social stimuli. The ROIs for this study were in areas whose activation was correlated with SPS under these conditions.

### 1.6 Study aims

We undertook this study to further validate and contribute to the broad understanding of SPS as an innate trait associated with a high level of perceptual attention. This trait accounts for a broadly-defined attentional survival strategy: high-level attention to detail. In the present research, we correlated HSP proxy questionnaire scores with DTI measures to assess any axonal microarchitecture measures associated with SPS, particularly in brain areas that might be related to primary and secondary perceptual processes. Importantly, we also wanted to determine if a subtle behavioral trait such as SPS could be detected using DTI. The results suggest that the heightened sensory processing in people with the SPS trait may be influenced by the anatomical microstructure of white matter in specific neocortical regions. Although previous fMRI studies had identified most of these general neocortical regions, the DTI-based results put a new focus on attention and flexibility, low-level primary sensory processing, empathy, emotion and depth of processing. Psychological trait characterization may benefit from diffusion tensor imaging methodology by identifying influential brain systems for the trait.

## 2 Methods

### 2.1 Participants

We used data from the Young Adult Human Connectome Project (YA-HCP) WU-Minn-Oxford consortium S500 release (Essen et al., 2012; Glasser et al., 2013), from which we used data of 408 subjects (243 females and 165 males) and self-report questionnaire data. Age of the subjects was between 22 and 36 years (mean age for females: 29.2 years, standard deviation: 3.4 years; mean age for males: 28.9 years, standard deviation: 3.6 years); ethnicity: 66.18 % White/European ancestry, 20.59% African-American, 7.84% Latino, 1.96% Asian or Nat. Hawaiian or other Pacific, 1.96% not reported, 1.47% more than one. The participants were free of documented psychiatric or neurological disorders.

### 2.2 Assessment of sensory processing sensitivity

The YA-HCP does not include the standard HSP scale for SPS, but does include multiple psychological measures. Therefore, we systematically identified and tested a subset of items in the HCP dataset that could serve as a proxy measure. First, the authors of the standard measure, the Highly Sensitive Person (HSP) Scale (Aron and Aron, 1997) examined the various self-report measures in the YA-HCP data, which are based on the NIH Toolbox, and selected 55 candidate items to assess SPS. We administered the 55 candidate items along with the standard HSP Scale (in counterbalanced order) to a sample of 401 mTurk workers (crowd sourcing marketplace for questionnaires; see Amazon Mechanical Turk, mturk.com). Of these 401, 19 failed one or more of four attention checks and 1 gave identical responses (all 1s or all 7s) to all the items in three of the main scales. The remaining sample of 381 included 174 women, 206 men, and 1 who did not indicate gender; mean age was 35.88 (SD = 11.23); 72% White/European ancestry, 8% African-American, 8% Asian, 7% Latino; 5% other.

We randomly divided the mTurk sample’s data into three groups, with the constraint of equal percentages of each gender in each group: Group 1, n=181, Groups 2 and 3, n=100 each. There were no significant differences in age or ethnicity between subgroups. In Group 1, we correlated each candidate item with the standard HSP Scale. Using those results, we explored several different subsets of the 55 items, checking each subset both for overall correlation with the HSP Scale and internal validity, and we identified a potentially optimal subset. Next, we administered this subset to Group 2 and made further adjustments, then tested this further adjusted version in Group 3 and also tested the reliability of this set of items in the overall HCP sample.

The resulting scale consisted of 17 items (see Table 1). In our mTurk sample of 381, the correlation of this 17-item measure with the standard HSP scale was 0.79; adjusting for reliabilities (0.93 for the HSP Scale, 0.79 for the 17-item proxy) yielded a deattenuated correlation of 0.89. This indicates that the 17-item subset is strongly parallel to the standard HSP Scale, and thus an appropriate measure of SPS. The alpha for these 17 items in the YA-HCP dataset was 0.62, which we considered marginally adequate, especially given that these items were taken from separate, not contiguous, scales in the YA-HCP dataset that measure using diverse response types. By contrast, in the mTurk sample the items from these scales were all administered close to each other. This only marginally adequate reliability does mildly undermine the strength of analyses, suggesting that some failures to find significant results may be due to the low reliability, although significant results obtained in spite of this are likely to be especially robust and may underestimate the actual effect size.

**Table 1:**
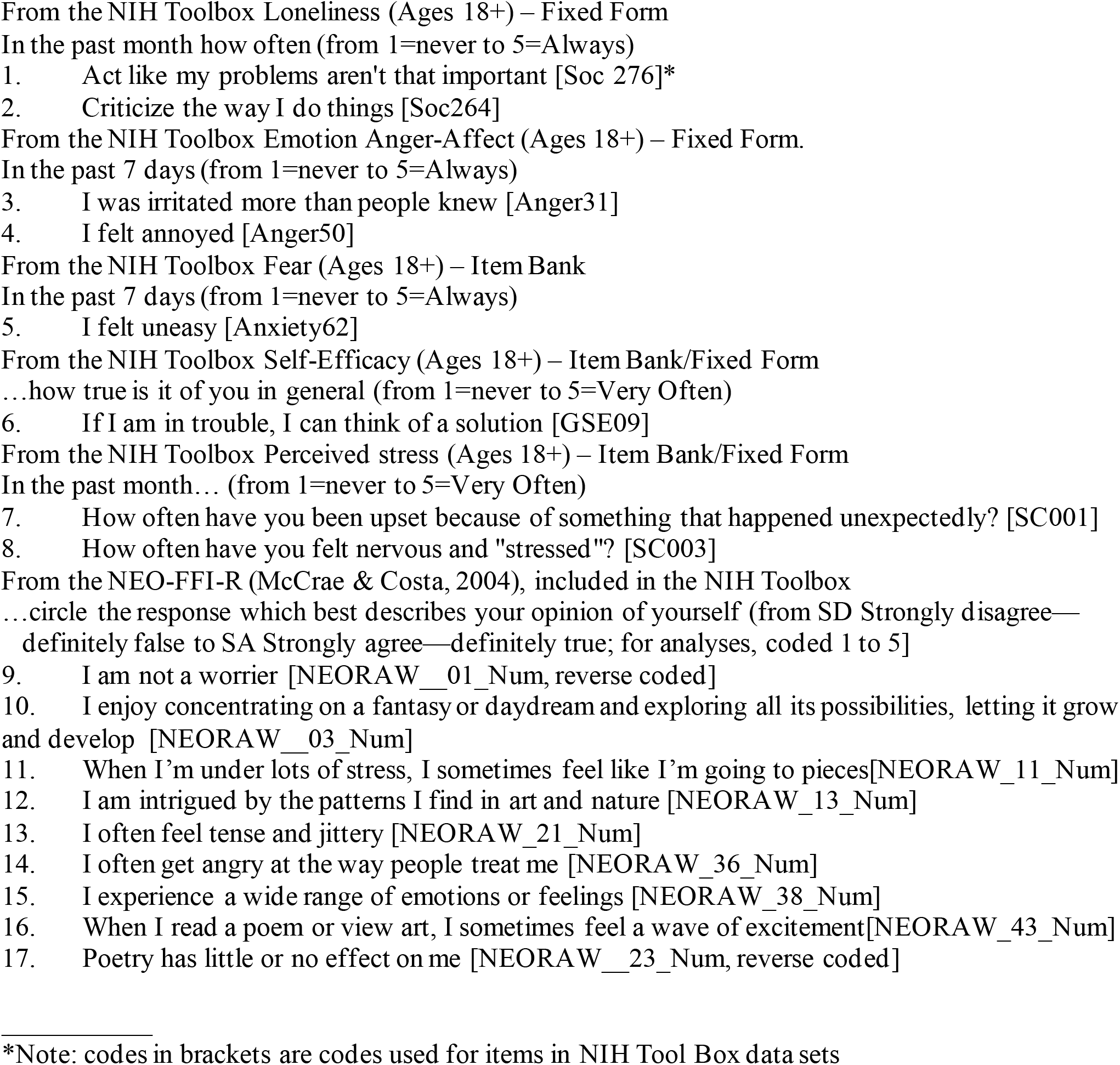
The HSP-Proxy Scale (17 items) Developed From Questions in the NIH Toolbox Used for the Human Connectome Project.

Finally, to control for negative affect, we further adjusted HSP scores. Typically, there is a substantial correlation between HSP scale scores and negative affectivity or neuroticism. Thus, it is standard practice to partial out scores on a measure of negative affect in SPS studies. We did so in the present study by creating standardized residuals of the proxy HSP Scale using mean NEO Neuroticism Scale scores from the sample. Thus, we computed neuroticism-score-adjusted studentized residuals, using a standard model in the statistical program SPSS, which shows HSP scores in terms of a standard deviation having negative and positive values. These scores were used to analyze the data and examples are shown in the graph in Figure 1.

**Figure 1.**
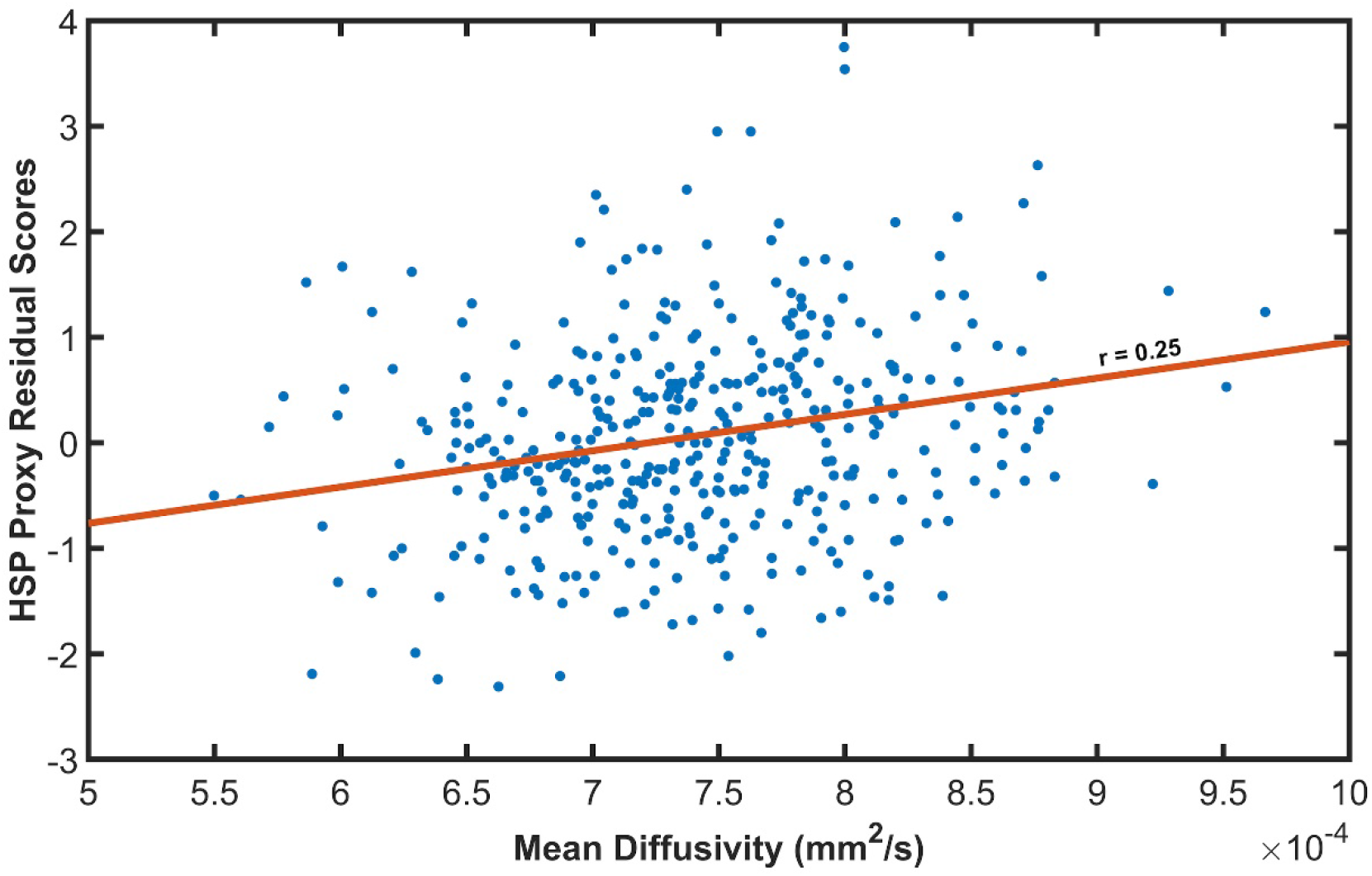
Mean diffusivity correlation with HSP residual score (standard deviations from the mean) in a voxel in the right medial prefrontal cortex, within the cingulum bundle (MNI: 12, 44, -10; see Fig. 2). This voxel showed the highest Cohen’s D effect size for the whole brain analysis (0.27; t=5.24).

### 2.3 Imaging data & processing

We used the minimally processed diffusion magnetic resonance images (dMRI) (Sotiropoulos et al., 2013), resulting in 90 diffusion-weighted images (DWIs) and 9 non-DWIs available for each participant with b-value of 1000 s/mm^2^. ExploreDTI version 4.8.6. (Leemans et al., 2009) and the REKINDLE (Tax et al., 2015) tensor estimation approach was used to calculate the voxelwise eigenvalues and eigenvectors. FA, MD, AD and RD maps were calculated from the fitted tensor model and warped to the MNI template. We also corrected for the gradient nonlinearities in the diffusion-weighted gradients (Bammer et al., 2003; Mesri et al., 2019), by using the voxelwise pattern of b-values and gradient direction during tensor estimation. The mean FA mask was calculated using all subjects and thresholded at FA > 0.2 to identify a white matter mask to limit the spatial extent of the statistical tests. Computations were performed on a Dell multi-core parallel processing system with 72 Intel Xeon E7-8870 v3 @2.10Ghz dual cores with 1 TB RAM.

### 2.4 Statistical tests

To investigate the correlation between the SPS scores and diffusion measures (FA, MD, AD, and RD), we used the nonparametric t-test via Permutation Analysis of Linear Models (PALM) (Eklund et al., 2016; Holmes et al., 1996; Nichols and Holmes, 2003; Winkler et al., 2014) with 10000 iterations. Significance was determined at p_corr_ < 0.05 using family-wise error rate (FWER) adjustment to correct for multiple comparisons, which corrected for multiple contrasts and modalities as well (Winkler et al., 2016b). Threshold-Free Cluster Enhancement (TFCE) (Smith and Nichols, 2009) was used to amplify p-values. Calculation speed was accelerated using the tail approximation (Winkler et al., 2016a). Because a large number of subjects can produce highly statistically significant results for small effects, Cohen’s D effect sizes were calculated and are the statistic we emphasize. FWER-corrected p-value maps were fed into the FSL tool automated atlas query (*autoaq*) to facilitate the anatomical interpretation of the statistically significant voxels.

In a separate analysis, the voxelwise effect size maps were thresholded at Cohen’s D > 0.1, regardless of the associated p-values, and the largest connected component (cluster) was selected using the *bwconncomp* MATLAB function. Exploring the non-trivial effect sizes may provide additional information regarding the spatial extent of the HSP scale – DTI metric relationship.

### 2.5 Region of Interest Analysis

In addition, we performed region of interest (ROI) analysis for primary sensory processing areas and areas associated with emotion processing based on previous data (Acevedo et al., 2017, 2014; Aron et al., 2010; Jagiellowicz et al., 2011). White matter ROIs were defined by the FreeSurfer (FS) ‘wmparc’ atlas. Thus, the ROIs used larger areas than those detected in previous fMRI studies, which may dilute any smaller regional significant effect. For example, previous studies found functional activations in the angular gyrus, temporoparietal junction and supramarginal gyrus that are all within the “inferior parietal” ROI region. Thus, we report the findings for some potentially diluted ROIs with P values >0.05, because these were planned comparisons and hypothesis-driven. The ROIs tested were white matter right and left: bankssts, caudal anterior cingulate, cuneus, entorhinal, fusiform, inferior parietal, inferior temporal, lateral occipital, lateral orbitofrontal, middle temporal, paracentral, parsopercularis, pericalcarine, postcentral, precuneus, superior parietal, transverse temporal (primary auditory), and insula.

ROI-based statistical testing was performed similarly to the voxelwise tests for all regional mean DTI metrics. PALM was utilized along with 10000 iterations with FWER adjustment and tail approximation. Furthermore, regional volume has been demonstrated to influence DTI estimates (Vos et al., 2011). Therefore, the ROI volume was considered as a co-variate of no-interest.

## 3. Results

### 3.1 Range of HSP Scores and diffusivity values

The raw HSP proxy questionnaire scores ranged from 1.71 to 3.82 (1 to 5, possible scores). HSP proxy residual scores, which are standard deviations from the mean, were used for the analyses and they ranged from -2.31 to +3.75 at the maximum effect size voxel (see Figure 1). The score range allowed an adequate sampling of HSP/SPS trait intensities. Based on approximate cutoffs from (Lionetti et al., 2018) latent class analysis of a large sample using the standard HSP scale, and adjusting their cutoff for mean and SD in the Proxy scale, a score of 2.97 or greater was considered to reflect the HSP trait as influential in everyday life. Sixteen percent (n=65) of our participants showed this range of scores, a population prevalence estimated by other studies of HSP. Thus, a relatively small but statistically adequate number of our participants would be considered a highly sensitive person, leading to small effect sizes. Mean diffusivity ranged from 5-10× 10^−4^ mm^2^/s, values found in normal, healthy brains (Lebel et al., 2008).

### 3.2 HSP proxy scores correlated with brain axon microarchitecture measures

#### 3.2.1 Whole brain, exploratory analysis

We found positive correlations between HSP-proxy scores and MD within the ventromedial cingulate, ventromedial and ventrolateral prefrontal cortex. The largest effects were in the right anterior/ventral subcallosal cingulum bundle, extending into the forceps minor of the corpus callosum (peak Cohen’s D = 0.269, P = 0.018). Fig. 1 shows the MD-HSP relationship at the peak effect size (MNI coordinates X:12, Y:44, Z: -10). Other MD effects were in the left anterior-ventral subcallosal cingulum bundle (Cohen’s D=0.243, p =0.028). Fig. 2/A visualizes the voxelwise results from all comparisons on the brain template, while Table 2 lists the regions, MNI coordinates, p-values and effect sizes.

**Table 2.** Whole brain exploratory analysis. Brain white matter areas showed positive and negative correlations between diffusion tensor imaging measures and proxy HSP scores for Sensory Processing Sensitivity. X/Y/Z denotes the MNI coordinates for the peak voxel within the cluster. Effect size is defined as Cohen’s D.

| Brain Region | x | y | z | # of Voxels | p-value | Peak effect size |
| --- | --- | --- | --- | --- | --- | --- |
| ***MD, Positive Correlation:***<br>*ventromedial prefrontal cortex/*<br>*Cingulate cortex:* |  |  |  |  |  |  |
| Anterior/Ventral Subcallosal<br>Cingulum Bundle and Forceps<br>minor of corpus callosum | 12 | 44 | -10 | 58 | 0.018 | 0.269 |
| Anterior/Ventral Cingulum<br>Bundle | -6 | 26 | -10 | 7 | 0.028 | 0.243 |
| ***RD, Positive Correlation:***<br>*ventromedial and lateral*<br>*prefrontal cortex* |  |  |  |  |  |  |
| Anterior/Ventral Subcallosal<br>Cingulum Bundle and Forceps<br>minor of corpus callosum | 10 | 36 | -12 | 29 | 0.041 | 0.170 |
| Anterior/Ventral Cingulum<br>Bundle | -6 | 26 | -10 | 3 | 0.033 | 0.228 |
| Anterior/Ventral Cingulum<br>Bundle | 18 | 48 | 0 | 21 | 0.032 | 0.232 |
| ***FA, Negative correlation;***<br>*motor/premotor cortical region* |  |  |  |  |  |  |
| Premotor area | 22 | -14 | 52 | 6 | 0.042 | -0.181 |

**Figure 2.**
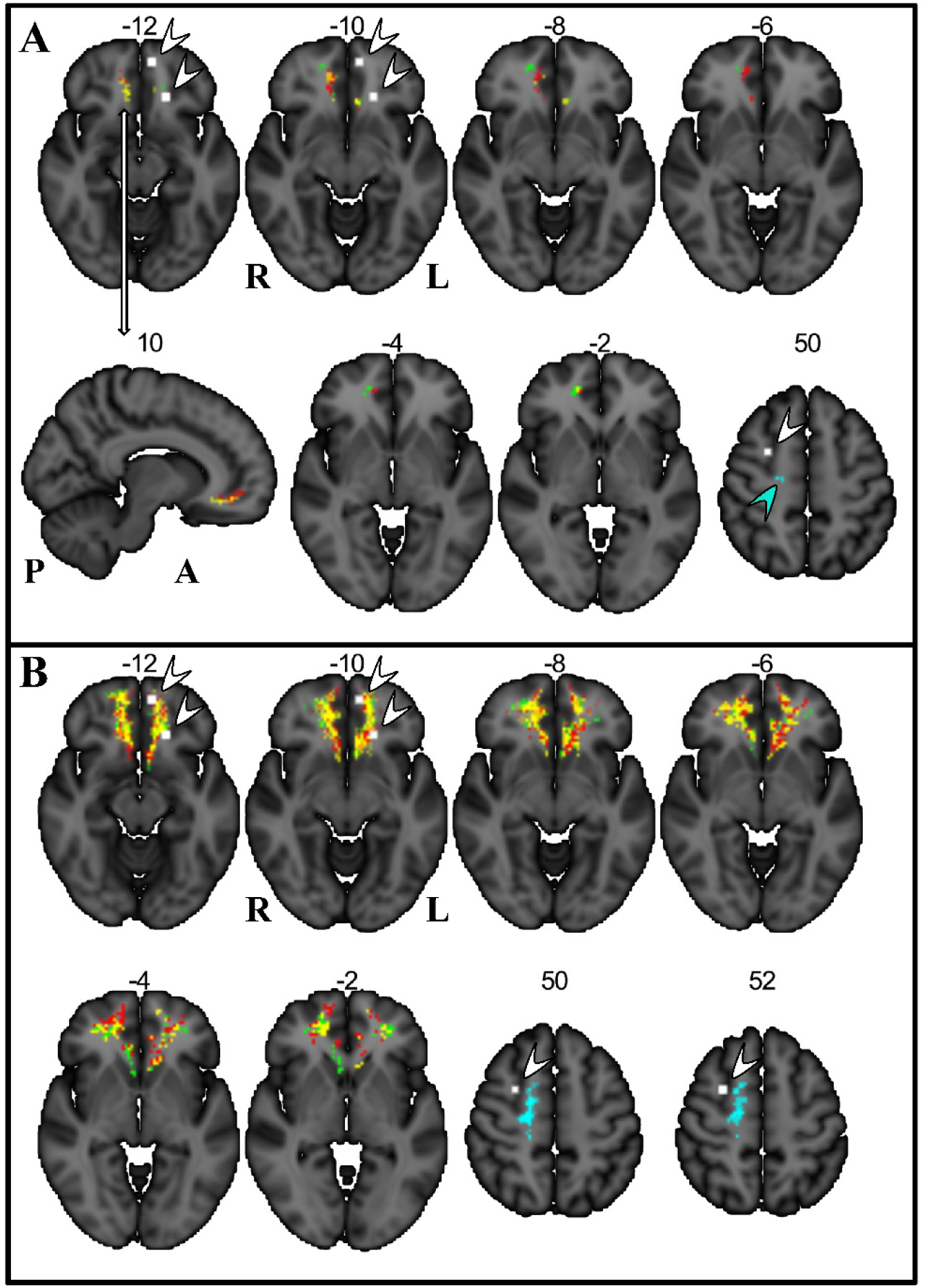
Voxelwise results presented in MNI stereotaxic space on axial and sagittal sections. A. Voxels in color show statistically significant correlations with HSP proxy questionnaire scores. They are located in the ventromedial prefrontal cortex, mostly on the right, within the forceps minor of the corpus callosum and the subcallosal cingulum bundle. The correlated voxels extend about 10 mm axially and about 25 mm antero-posteriorly. Significant voxels are also on the left, mostly in the subcallosal area. B. Colors show largest voxel clusters above Cohen’s D effect size of 0.10 without considering statistical significance (see Methods). Red: positive correlation with MD (mean diffusivity); Green: positive correlation with RD (radial diffusivity); Yellow: the overlap of MD and RD; Blue and blue arrow: negative correlation with FA (fractional anisotropy); White voxels and white arrows: locations of fMRI activations that correlated with HSP questionnaire scores in a previous study when participants viewed a romantic partner and a stranger, happy or sad (Acevedo et al., 2014).

There was also a positive correlation for RD in regions overlapping the MD effects in the right anterior/ventral cingulum bundle and forceps minor of the corpus callosum (Cohen’s D=0.170; p=0.041) and in a second region of the cingulum bundle (Left, Cohen’s D=0.228; p=0.033 Right, Cohen’s D=0.232; p=0.032; Table 2; Fig. 2/A).

Connected voxels showed a continuous band in the ventral cingulate and ventromedial prefrontal white matter from the posterior genu of the corpus callosum and radiation of the straight gyrus to the frontal pole, including the superior longitudinal fasciculus, inferior fronto-occipital fasciculus, uncinate and arcuate fasciculus as shown in Fig. 2/B. Furthermore, these results extended into the ventrolateral prefrontal cortex. White matter anatomical findings were near gray matter functional activations found previously in SPS subjects reacting to emotional stimuli (Acevedo et al., 2017) and to a romantic partner’s emotional facial expression as shown in (Acevedo et al., 2014).

We also found a negative correlation between HSP-proxy scores and FA (Cohen’s D=-0.181; p=0.042) in the right premotor cortex area in the region of the origin of the corticospinal tract (Fig 2/A). Connected voxels covered a large area that included white matter in the region of the dorsomedial prefrontal cortex, premotor cortex, precentral gyrus (motor cortex), post central gyrus (somatosensory cortex), and supramarginal gyrus (somatosensory association cortex) in the parietal lobe (Fig. 2/B, 3). Fig. 3 shows the 3D render of the FA cluster, after thresholding only for effect size. This cluster was near gray matter functional activations observed previously in SPS subjects reacting to a romantic partner’s emotional facial expression as shown in Acevedo et al. (2014) (Acevedo et al., 2014) (Fig. 2A/B).

**Figure 3.**
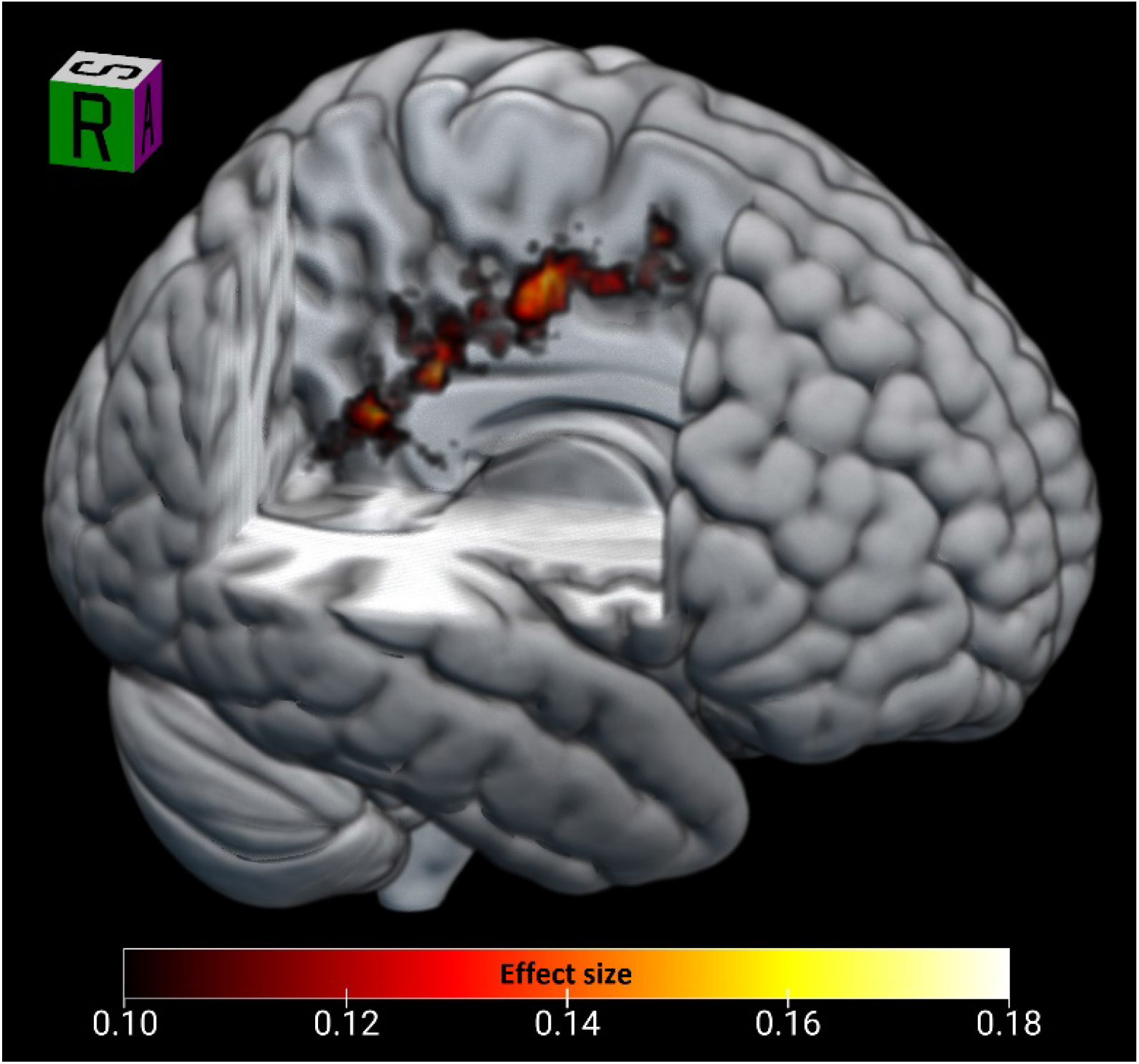
3D render of the largest FA cluster (negatively correlated with HSP score). The cluster extends from the premotor cortex through the primary motor and somatosensory cortex into the supramarginal gyrus.

#### 3.2.2 Region of interest analysis

Region of interest analyses showed statistically significant positive RD and negative FA correlations with neuroticism-adjusted HSP-proxy scores in two areas. Figs. 4 and 5 show the ROI analysis results, while Table 3 shows the numerical summary. RD was positively correlated with scores in the left precuneus (Cohen’s D=0.1509, p=0.04; Fig. 4/C) and marginally on the right (Cohen’s D=0.143, p=0.06, Fig. 4 B/C). FA was negatively correlated with scores in the right parsopercularis/inferior frontal gyrus (Cohen’s D=-0.165, p=0.02).

**Table 3.**
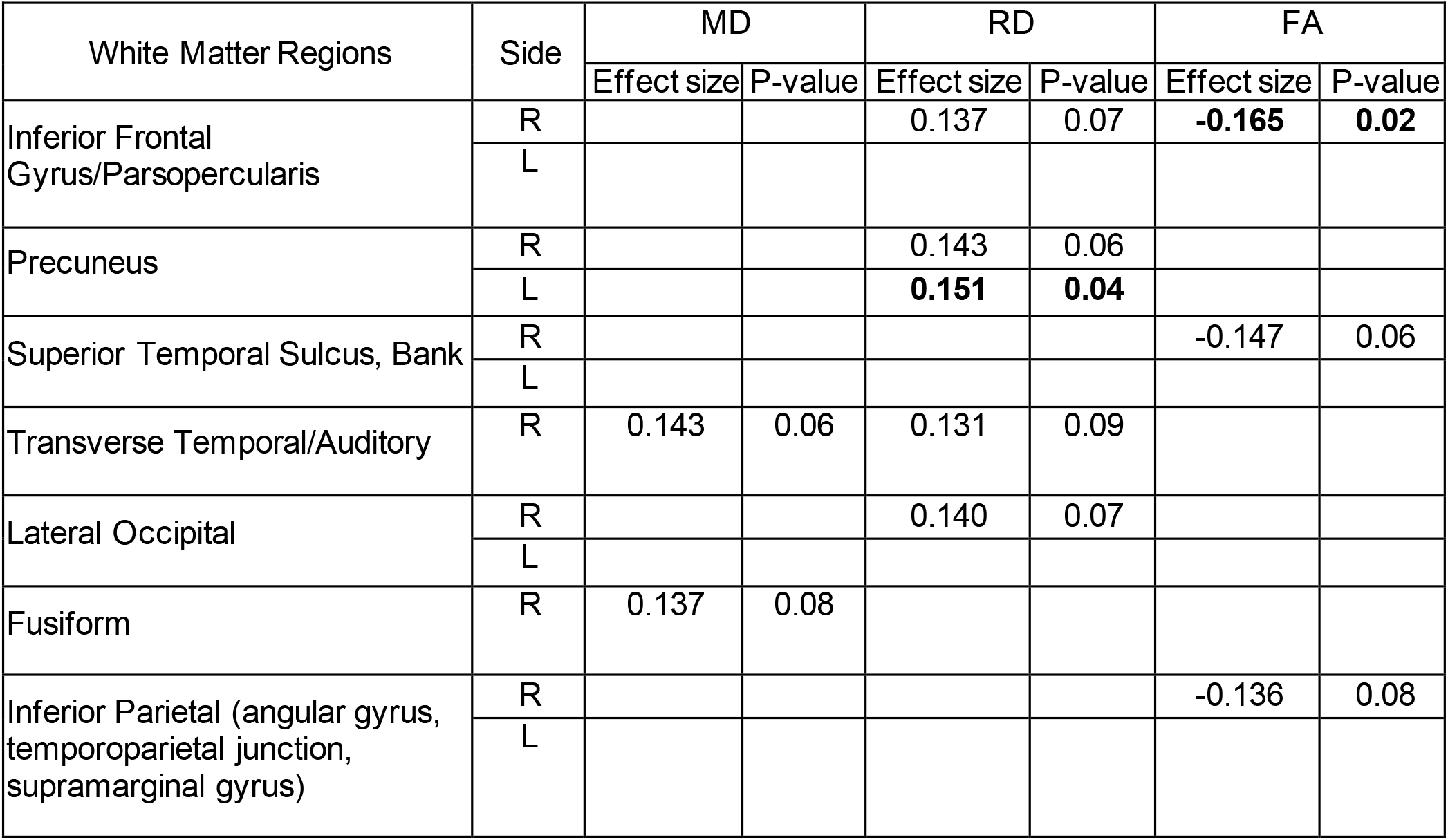
Region of Interest analysis. White matter areas with Cohen Effect Sizes >0.1 and p values at statistically significant, or marginally significant values.

**Figure 4.**
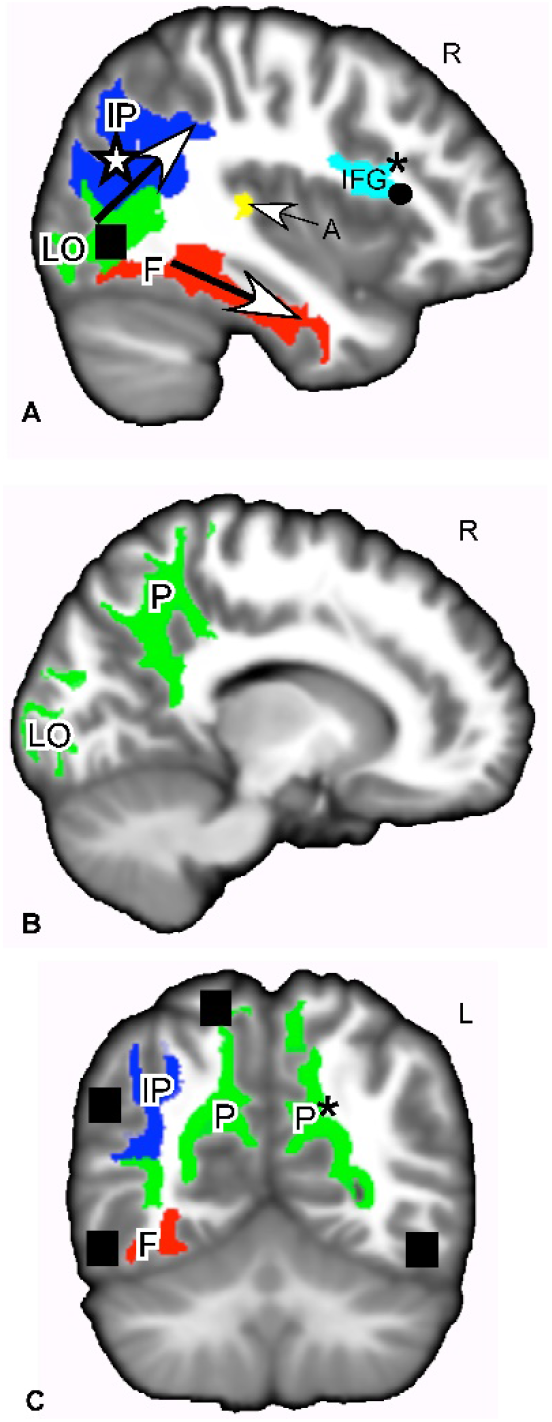
White matter regions of interest where MD, RD or FA showed positive or negative correlations with HSP proxy questionnaire scores. A. Sagittal slice through ROIs that showed a correlation between HSP-proxy scores and DTI measures. These ROIs are white matter of the ventral pathway for visual processing (LO,F bottom arrow), dorsal pathway for visual processing (IP, top arrow), primary auditory processing (A), and empathic responses (IFG). B. Sagittal slice through regions that showed a positive correlation between RD and the HSP-proxy scores. The ROIs are white matter of the ventral and dorsal visual pathways (LO, P). C. Coronal slice through white matter regions involved in the ventral and dorsal visual pathway (F,IP,P) that showed correlations between questionnaire scores and MD, RD and FA. These regions are connected to gray matter areas associated with SPS (▮) Red: MD positive correlation; Green: RD positive correlation; Yellow: MD+RD positive correlations; Dark Blue; FA negative correlation; Light Blue: FA negative+RD positive correlation. *p<0.05. Otherwise, p values ranged from 0.06-0.09. Effect sizes -0.165 to + 0.151. See Table 3. Shape symbols indicate where previous fMRI and behavioral studies of SPS found activations implicating heightened sensory processing, empathy, attention and self-other processing. ▮Jagiellowicz et al., 2010 ☆Acevedo et al., 2014, 2017 • Acevedo et al., 2014 Abbreviations: A, primary auditory cortex. F, fusiform. IFG, inferior frontal gyrus. IP= Inferior parietal/angular gyrus/temporoparietal junction/supramarginal gyrus. LO, lateral occipital area. P, precuneus. MD, mean diffusivity. RD, radial diffusivity. FA, fractional anisotropy.

**Figure 5.**
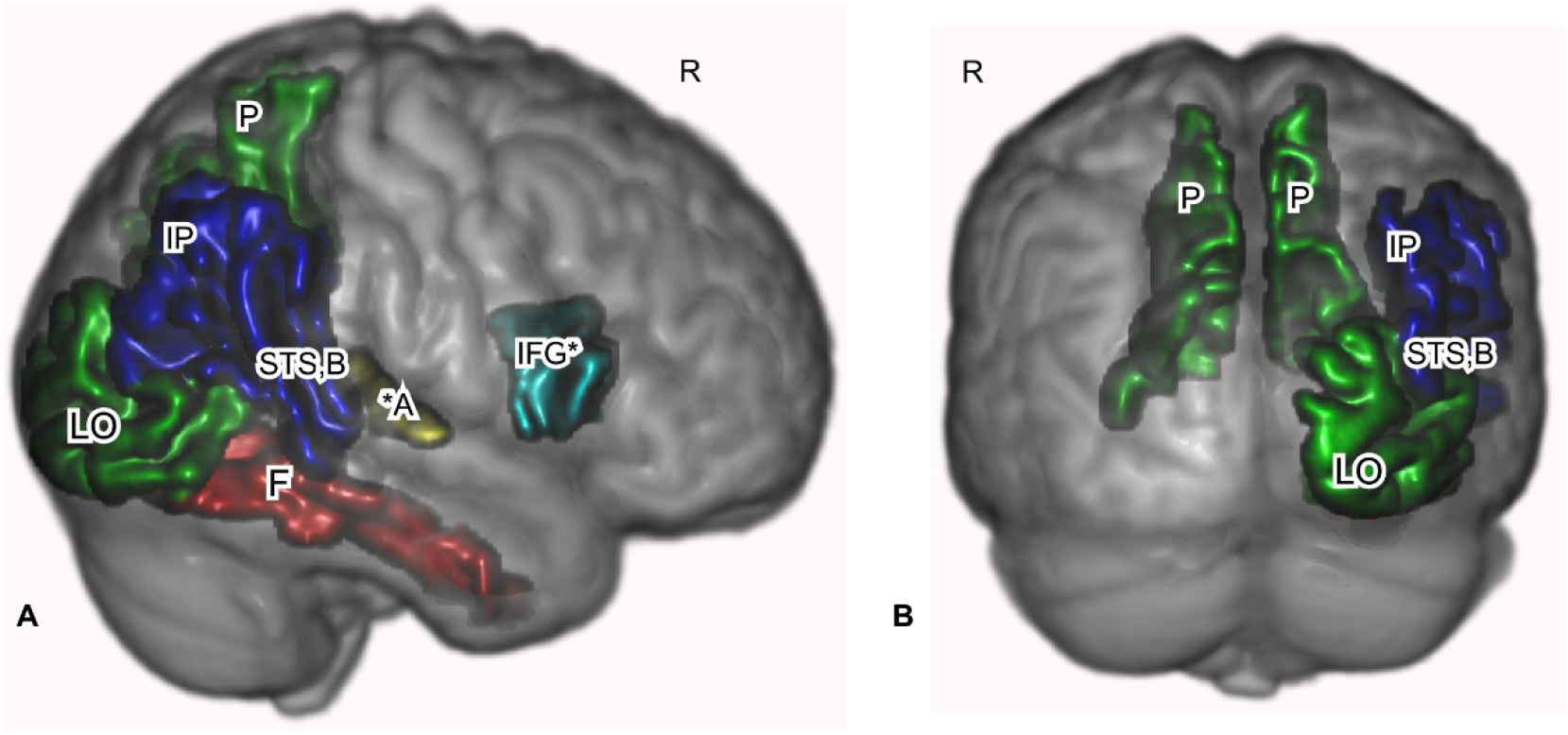
Primary sensory, higher order sensory processing and cognitive processing regions were identified as correlated with the HSP proxy scores. Freesurfer-based ROIs are shown for white matter where Cohen’s D effect size was -0.147 to +0.151, rendered in 3D on a template brain. Areas include the general path of the dorsal and ventral visual pathways. A. Sagittal view of the right side. B. Posterior view. Red: MD positive correlation with HSP proxy questionnaire scores. Green: RD positive correlation. Blue: FA negative correlation. Gold: MD and RD positive correlation. Aquamarine in IFG: RD and FA correlations. *p<0.05. Otherwise, p values were 0.06-0.09. See Table 3. Abbreviations: A, primary auditory cortex. F, fusiform. IFG, inferior frontal gyrus. IP, inferior parietal including angular gyrus/temporoparietal junction/supramarginal gyrus. LO, lateral occipital cortex. P, precuneus. STS,B-bank of the superior temporal sulcus. MD, mean diffusivity. RD, radial diffusivity. FA, fractional anisotropy.

We report other areas at p>0.05, because they are primary visual sensory or higher order sensory processing areas that were activated in previous fMRI studies: the primary visual cortex (right lateral occipital area, RD: Cohen’s D=0.14, p=0.07, see Table 3; Fig. 4A/B), the right fusiform gyrus (MD; Cohen’s D=0.137, p=0.08, see Table 3; Fig. 4A/C), the right bank of the superior temporal gyrus (FA; Cohen’s D=-0.147, p=0.06; see Table 3; Fig. 4/A), and the inferior parietal cortex that includes the angular gyrus, temporoparietal junction and supramarginal gyrus (FA Cohen’s D: -0.136, p=0.08; Fig. 4/A). These areas (Fig. 4/A) are part of the “ventral visual stream” that identifies “what” in the visual field and the “dorsal visual stream” that identifies “where” in the visual field (Milner and Goodale, 2008; Ungerleider and Haxby, 1994). Functional activations of these regions were seen in SPS individuals during a visual discrimination task (Acevedo et al., 2017, 2012; Aron et al., 2010; Jagiellowicz et al., 2011). Finally, MD was positively correlated with HSP-proxy scores in the right transverse temporal/primary auditory cortex (Cohen’s D=0.143, p=0.06; Fig. 4/A) and on the left (Cohen’s D=0.131, p=0.09).

## 4. Discussion

### 4.1 Overview

This study established an anatomical correlate in the white matter of the brain for SPS individuals: those with the highest neuroticism-adjusted HSP proxy scores showed the greatest RD, FA and MD effects in several neocortical areas compared to those with the lowest HSP scores (not highly sensitive). Whole-brain, exploratory effects were greatest in brain regions involved in (a) higher-order emotion and reward processing: the ventromedial prefrontal cortex; and (b) in regions involved in empathy, self-other processing, attention and flexible coding of the environment: the premotor cortex and supramarginal gyrus. Compared to whole brain results, smaller ROI effects were also seen in self-other processing areas (IFG, precuneus, fusiform, angular gyrus), higher order visual processing regions of the ventral and dorsal pathways (precuneus, inferior parietal, temporoparietal junction, STSb), and primary sensory processing areas such as the lateral occipital and transverse temporal (primary) auditory cortex.

All of these results are consistent with behavioral observations for SPS, which include sensory sensitivity, a tendency to be overwhelmed by sensory stimuli, and more attention to emotional and visual details of stimuli than others who do not show these traits (Aron et al., 2012; Aron and Aron, 1997). The localization of the higher-order visual processing effects is also consistent with the previous studies, as reviewed in the Introduction, that used functional MRI to assess details of visual scenes and emotional reactions in highly sensitive people (Acevedo et al., 2017, 2014; Jagiellowicz et al., 2011). Thus, regional *functional* effects previously described were confirmed by microstructural effects. In addition, this is the first study to suggest involvement of primary sensory processing areas in the cortex, such as the visual and auditory cortex. The study also highlights the premotor cortex, with its connections to the supramarginal gyrus and attention functions. Somatosensory as well as environmental stimuli like vision are new possible highlights. Further, the study suggests a novel focus on the functions of the ventromedial prefrontal cortex. Finally, this is one of few studies to show small DTI anatomical effects within axonal tracts for normal-range behavioral traits, in this case sensory processing. Other studies have investigated neurologically normal subjects for DTI effects, mostly using the “big five” personality traits (Xu and Potenza, 2012), but these traits overlap very little with the HSP Scale, other than with neuroticism (which is controlled for in this study). In this study of a normal trait, the results indicate novel behavior-related brain regions to explore in future studies.

### 4.2 Ventromedial prefrontal cortex

The largest effects were in white matter of the ventromedial and ventrolateral prefrontal cortex (De La Vega et al., 2016), on both right and left sides, but with more voxels on the right. These tracts connect major limbic system components: the hippocampus/peri-hippocampal cortex and amygdala to the medial prefrontal cortex. The subcallosal cingulum area, functionally connected with the ventromedial and ventrolateral prefrontal cortex (Dunlop et al., 2017) is known for its influence on mood, especially depression (Dunlop et al., 2017; Mayberg et al., 2013). The cingulum is also structurally connected to several of the ROIs that were correlated with HSP-proxy scores in this study: the precuneus, bank of the superior temporal sulcus, and supramarginal gyrus (Bathelt et al., 2019).

The ventromedial and lateral prefrontal cortex areas are described by (Hiser and Koenigs, 2018) as having three broad domains of psychological function: Decision making based on reward and value (Sescousse et al., 2013); generation and regulation of negative emotions; and, social cognition, such as facial recognition, theory of mind, and processing self-relevant information. These of course interact strongly, especially given the social nature of humans, as in the function of the vmPFC in making moral decisions (Cameron et al., 2018) or in the “intuitive feeling of rightness” that guides decision making, often social in nature, as a function of memory retrieval (Hebscher and Gilboa, 2016).

The major results in the vmPFC in this study, as well as in the preceding fMRI studies (Acevedo et al., 2017, 2014), point to perhaps the most important aspect of SPS, which is “depth of processing” (Aron et al., 2012). The term is based on cognitive conceptualizations of levels or processing, with the idea that processing to deeper levels with more detailed cognitive contexts leads to better memory and better learning overall (Leow, 2018; Lockhart et al., 1976). The hypothesized evolutionary development of depth of processing in SPS is based on a computer simulation demonstrating that unusual responsivity to the environment will evolve when there are enough payoffs for an individual difference in noticing details, as long as most individuals do not notice these details (Wolf et al., 2008). (If all individuals did, there would be no special benefit.) That is, SPS is considered fundamentally an individual difference in “depth of processing” through careful observation of situation/time A in order to compare those details in memory to situation/time B and gain any potential benefits others miss. Individuals without the trait are thought not to process A as carefully, a strategy which is often equally or more effective, since B may bear no resemblance to A or there may be little reward in noticing any resemblances.

This type of careful processing relies on emotional motivation, the desire for rewards, such as winning, and the desire to avoid fear-related stimuli, such as losing (Baumeister et al., 2007). Hence understanding SPS as fundamentally about depth of processing for decision making based on reward value and social value is consistent with the significant differences that were found in the vmPFC, with its close connections to emotion-related memory processing areas such as the hippocampus and amygdala. Memory enhanced by motivation is, again, key to SPS, and specific activations occur for those high in SPS when processing emotional stimuli. For example, Acevedo et al (Acevedo et al., 2017) found in their comparison of responses to positive and negative stimuli that there were considerable differences in regional brain activation for high and low SPS in the vmPFC, in the same areas where microstructural differences were found in this study (Fig. 2), and in regions that mediate memory, attention, awareness, and reflective thinking.

#### Theory of Mind

In a meta-analytic review of the Theory of Mind, (Mar, 2011) highlighted a core mentalizing network: the mPFC, precuneus, bilateral pSTS, bilateral angular gyri, and the right IFG. These structures showed microstructural differences associated with SPS in this study, notably the mPFC, right IFG, precuneus, the bank of the STS and angular gyrus region. Second, this mentalizing network overlaps with the narrative comprehension network in a number of areas, including the mPFC, bilateral pSTS/TPJ, precuneus, and possibly the right IFG, again areas implicated by this study. Theory of Mind is central to the core concept of SPS, in that this survival strategy would require more reflection on another’s behavior in order to accurately see from their perspective and correctly attribute to them their motivations and intentions.

### 4.3 Premotor cortex, attention and somatosensory processing

Finally, regarding the extensive cluster from the left premotor cortex to the post-central somatosensory and supramarginal gyrus, it could be expected that the arc of axonal effects we found in sensory processing areas from posterior to anterior (Figs. 2B, 4A and 5A) would include the premotor cortex, where the hypothesized deep processing associated with SPS would result in the preparation for action. This potential somatosensory/posterior parietal/premotor cortex involvement in SPS is a novel contribution to its understanding and may be helpful in future studies.

The importance of this broad premotor area continues to evolve (Rizzolatti et al., 1987). Rizzolatti et al. suggested a theory of attention focused on this area, presenting evidence that the premotor areas were the source of attention rather than a separate attention-directing mechanism. That is, attention is turned to a stimulus within the premotor area, so that attention consists of nothing more than preparation for a motor activity (e.g., an eye movement towards a stimulus deemed important or auditory areas preparing for a sound). Although refinements and extensions (Wollenberg et al., 2018) have occurred, this view of attention is still tenable, according to experiments by (Schubotz and Von Cramon, 2003).

Schubotz and Cramon distinguished the left premotor cortex, correlated in our study with SPS, as associated with nonspatial tasks and rapid acquisition of new motor sequences. Overall, evidence regarding the premotor area suggests “environmental features do not have to remind us of specific actions or movements to induce premotor activation on a more or less conscious level. Rather, features are represented in a highly fragmented format that allows for instant recombination and very flexible coding of any currently attended environment” (p. 126). (Schubotz, 2007) presented considerable evidence that the premotor area (along with most of the areas in the brain associated with SPS) helps in the prediction of events.

Further, a mean diffusivity study by (Takeuchi et al., 2019) found that an area including the premotor cortex plays a major role in emotional salience and empathy. This area was also activated in the (Acevedo et al., 2014) fMRI study finding empathy for happy and distressed partners and strangers.

Attention, flexible coding, prediction, somatosensory processing and empathy all fit with the theory that SPS involves attending to subtle stimuli that may be relevant for survival and predicting those environmental details that what will be relevant in future environments. Overall, these whole brain, exploratory analyses provide a picture that is consistent with behavior associated with SPS.

### 4.4 ROI results

The particular ROIs with clear statistical significance were the left precuneus (RD, Cohen’s D = +0.151, p=0.04), part of the dorsal visual stream, and the right parsopercularis/IFG (FA, Cohen’s D = 0.165, p=.02), associated with empathy. For the precuneus, the other side was marginally significant (p=0.06), while for the IFG, the same side was marginally significant for RD as well (Cohen’s D = 0.137, p=0.07). Also, the transverse temporal gyrus/primary auditory cortex (A1), part of the dorsal auditory stream was marginally statistically significant (MD, Cohen’s D 0.143, p=0.06).

As for the precuneus, fMRI studies (Cavanna and Trimble, 2006) of healthy subjects suggest it plays a major role in visuo-spatial imagery, episodic memory retrieval, and self-processing tasks, such as the experience of agency and taking the first-person perspective. All of these activities are more prominent in those high in SPS, and the precuneus was another area often correlated with SPS in fMRI studies. There is also an hypothesized role for the precuneus in consciousness itself (Cavanna, 2007), along with areas nearby in the posteromedial parietal cortex. It is especially active during the conscious resting state, but is deactivated when consciousness decreases (e.g., sleep, anesthesia, Alzheimer’s). Indeed, it has been proposed that it is part of a larger network that correlates with self-consciousness, as it engages in self-related mental representations, self-reflection, and autobiographical memory retrieval. Meditation, which creates states of restful alertness, is associated with microstructural differences in the precuneus in practitioners compared to controls (Avvenuti et al., 2020; Shao et al., 2016). Without suggesting that SPS somehow results in more consciousness, it may well demonstrate an internal tendency for more awareness and integration of diverse aspects of inner and outer experience. Although the insula was not a factor in this study, it has a similar role in the brain and was found to be more active in the fMRI studies of SPS already cited, and has also sometimes been described as the “seat of consciousness” (Craig, 2009).

The right parsopercularis/IFG region was negatively associated with HSP proxy scores and FA (p=0.02), and positively associated with RD (p=0.07). IFG functions may be particularly important to recognize in further studies. In an fMRI study, the IFG region was positively associated with HSP scores during positive emotion conditions while looking at a spouse or stranger (Acevedo et al., 2014). It has been identified with a mirror neuron system (Iacoboni et al., 1999; Jabbi and Keysers, 2008; Van Overwalle and Baetens, 2009) that responds to the movements of others, and may facilitate the understanding of others’ intentions and feeling of empathy. We previously suggested that this system’s activation is consistent with HSPs’ bias toward noticing positive expressions in others and high empathy (Acevedo et al., 2014).

Some of the ROI effects are along the dorsal and ventral visual/auditory pathways, which is especially noteworthy as they are, again, what one would expect given that the hypothesized fundamental characteristics of the SPS trait are depth of processing of sensory experiences and awareness of subtleties. These pathways are described as identifying the “what” (ventral stream) and “where” (dorsal stream) of what is seen and heard, taking sensory experience beyond its initial input (Milner and Goodale, 2008). The ventral stream in particular is associated with object recognition and form representation, the “what,” and is strongly connected to the medial temporal lobe, which stores long term memories; the limbic system, which controls emotions; and joins with the dorsal stream, which identifies the “where.” The dorsal stream is said to guide actions and recognize where objects are in space. It stretches from the primary visual cortex in the occipital lobe into the parietal lobe. It contains a detailed map of the visual field and serves to detect and analyze movements. Thus, it commences with purely visual functions, ending with spatial awareness at its termination. As with the ventral stream, processing of sensory input along the dorsal stream becomes “deeper” or more elaborate. It ends up contributing to recognizing spatial relations, body image, and physical coordination. Again, as SPS has been described, the trait is not characterized by better initial sensory perception, better hearing or eyesight, but by more complete processing of what is perceived along these two visual/auditory pathways. However, the potential involvement of primary visual, auditory and somatosensory areas suggested in this study leads to other questions for study of the most basic sensory detection and discrimination functions of these areas that may impact the higher-order processing regions.

The A1 cortex that we included as a ROI for this study is thought to operate very early in the recognition of sounds. For example, a study by (Warrier et al., 2009) found that non-Mandarin-speaking subjects who could successfully form an association between Mandarin Chinese “pitch patterns” and word meaning were found to have transverse temporal gyri (A1) with larger volume than subjects who had difficulty learning these associations. Successful completion of the task also was associated with a greater concentration of white matter in the left A1 of the subject. In general, larger transverse temporal gyri seemed to be associated with more efficient processing of speech-related cues, which could aid the learning and perceiving of new speech sounds. The A1 cortex is also associated with inner speech, what (Hurlburt et al., 2016) might be considered a more advanced level of processing, but still preceding speech production. Although to date there are no studies of auditory functioning associated with SPS, it would seem to be a fruitful area for future research.

The superior temporal sulcus (STS bank, p=0.06) is seen primarily as an area for higher visual processing. (Hein and Knight, 2008), in a review of carefully selected fMRI studies, concluded that the majority of findings implicate the STS in broader tasks involving theory of mind, audiovisual integration, motion processing, speech processing, and face processing. They conclude that rather than trying to pinpoint where in the STS these occur, it is best to view the function of the STS as varying according to the nature of network coactivations with different regions in the frontal cortex and medial temporal lobe during a particular task. This view is more in keeping with the notion that the same brain region can support different cognitive operations depending on task-dependent network connections, emphasizing the important role of network connectivity analysis in neuroimaging. It is consistent with current hypotheses about SPS that those high in SPS would show greater microstructural differences in an area associated with diverse types of processing (motion, speech, face, audiovisual) as well as theory of mind.

### 4.5 Microstructural differences: the physiological impact of positive MD/RA and negative FA correlations

The physiological impact of positive MD/RA and negative FA correlations is unclear (Soares et al., 2013). The findings in this study are smaller than those seen in previous studies of disease progression or aging (Nir et al., 2013; Voineskos et al., 2012). There is no neurological or behavioral pathology in the group that we studied. The psychological traits measured are subtle and part of the normal range of human behavior. Thus, the effects are part of a normal variability in the population, but may be markers of slight anatomical differences in axon size and organization, perhaps impacting the speed of communication among regions (Horowitz et al., 2015).

This study joins others that have looked at microstructural correlates of normal individual differences, such as personality traits (Xu and Potenza, 2012) and cognitive abilities (Bathelt et al., 2019). Since such phenotypic differences can be caused by multiple genetic and environmental effects, looking for common microstructure may be another useful way to identify such differences.

### 4.6 Limitations

A limitation of the study is its reliance on a proxy measure of SPS. Few questions in the YA-HCP questionnaire dataset addressed sensory processing directly. However, the proxy measure was found in independent samples to have a strong correlation with the standard measure (r=0.79; r=0.89 adjusting for reliabilities). Future aims of research more generally will be to replicate these findings and to examine the relation of SPS to DTI in younger and older age groups and in other populations.

## 5. Conclusions

This is the first study to investigate the relation of SPS to neural anatomical measures using DTI. The study employed a relatively large sample of young healthy individuals, and it has identified several brain systems that may be critical to fully understanding the SPS trait, such as parietal/premotor connections. The study has also confirmed the involvement of several brain systems and areas previously correlated with SPS in fMRI studies, such as those for empathy and higher-order visual scene processing. Future research should focus on primary visual and auditory processing; higher-order somatosensory processing; attention flexibility and reward value processing. Also, the development of a proxy measure allows future research to examine the relation of SPS to other variables in the large YA-HCP sample, such as various genetic, functional imaging, and self-report data. Finally, DTI may be a valuable and fairly straightforward approach for future psychological and anatomical studies of normal individual differences because the scan can be acquired quickly and is often included in the usual battery of clinical scans.

## 8. Data statement

All data are available at https://db.humanconnectome.org.

